# *In vitro* activity of imipenem/relebactam alone and in combination against cystic fibrosis isolates of *Mycobacterium abscessus*

**DOI:** 10.1101/2024.12.10.627836

**Authors:** Madeline Sanders, Sun Woo Kim, Aditi Shinde, Danielle Fletcher-Williams, Eric Quach, Paul Beringer

## Abstract

**Background:** *Mycobacterium abscessus* complex (MABSC) is an opportunistic pathogen that causes chronic, difficult-to-treat pulmonary infections, particularly in people with cystic fibrosis (PwCF), leading to rapid lung function decline, and increased morbidity and mortality. Treatment is particularly challenging due to the pathogen’s resistance mechanisms and the need for prolonged multidrug therapy, which is characterized by poor clinical outcomes and highlights the urgent need for novel therapeutic strategies. Imipenem/relebactam, a novel β-lactam-β-lactamase inhibitor combination, demonstrates *in vitro* activity against resistant MABSC strains and effective pulmonary penetration. Prior literature indicates synergistic activity of imipenem with various antibiotics against *M. abscessus*.

**Objectives:** This study aims to evaluate the *in vitro* activity of imipenem/relebactam, alone and in combination with various antibiotics, against MABSC clinical isolates from PwCF (*n*=28).

**Methods:** Susceptibility and synergy were assessed using broth microdilution and checkerboard assays. Extracellular time-kill assays were performed to evaluate the bactericidal activity of synergistic three-drug combinations containing imipenem/relebactam.

**Results:** Imipenem/relebactam demonstrated potent *in vitro* activity against clinical MABSC isolates, exhibiting substantial synergy with cefuroxime, cefdinir, amoxicillin, and cefoxitin. Rifabutin, azithromycin, moxifloxacin, clofazimine, and minocycline also demonstrated additive effects with imipenem/relebactam. Extracellular time-kill assays identified imipenem/relebactam+cefoxitin+rifabutin and imipenem/relebactam+cefoxitin+moxifloxacin as the most effective combinations.

**Conclusions:** These findings suggest that imipenem/relebactam may offer a significant advancement in the management of MABSC infections in PwCF. The promising efficacy of multidrug regimens combining imipenem/relebactam with agents like cefoxitin, azithromycin, moxifloxacin, clofazimine, and rifabutin highlights potential therapeutic strategies.

## 1. Introduction

Cystic fibrosis (CF) is an autosomal recessive, multisystem disorder caused by mutations in the cystic fibrosis transmembrane conductance regulator (CFTR) gene, which disrupts ion and water transport across epithelial cells [1,2]. This dysfunction results in increased mucus viscosity and impaired secretion, particularly in the respiratory and gastrointestinal tracts. The thickened mucus impairs mucociliary clearance and depletes airway surface liquid volume, triggering damaging cycles of airway obstruction, inflammation, and infection that progressively impair lung function, ultimately leading to respiratory failure [2]. As lung function deteriorates, people with CF (PwCF) become increasingly vulnerable to chronic bacterial infections.

Nontuberculous mycobacteria (NTM) are increasingly being isolated from the sputum of PwCF, with prevalence estimates rising from 1.3% in 1984 to 10.1% in 2023 [3–5]. The prevalence of NTM increases with age, from an average of 10% in children aged 10 years to over 30% in adults above 40 [3]. Additionally, between 2010 and 2019, the average annual incidence of NTM pulmonary infections among PwCF in the U.S. was 58.0 cases per 1,000 individuals, with a significant annual increase of 3.5% [6]. Over time, NTM can cause progressive inflammatory lung damage, a condition termed “NTM pulmonary disease” (NTM-PD). PwCF are predisposed to NTM-PD in part due to reduced CFTR-mediated reactive oxygen species generation within macrophages, leading to reduced intracellular killing [7]. Prior literature has also shown that NTM have evolved mechanisms to evade host immune responses and survive intracellularly [8–9], presenting significant challenges for antibiotics to effectively target and eradicate the infection. Prevalence surveys worldwide show that the slow-growing *Mycobacterium avium* complex and rapidly-growing *Mycobacterium abscessus* complex (MABSC) account for over 95% of NTM lung disease cases in PwCF [10]. Recent epidemiological studies have demonstrated that the presence of MABSC in the airways of PwCF is associated with more rapid lung function decline and higher mortality [3]. Furthermore, a large-scale longitudinal study identified *M. abscessus* as the most significant contributor to lung function deterioration among NTM and gram-negative bacteria in this population [11].

Treatment of MABSC pulmonary disease (MABSC-PD) remains highly challenging and necessitates prolonged therapy with multiple antibiotics. Current treatment guidelines include an initial intensive phase consisting of oral azithromycin in combination with several intravenous (IV) antibiotics (e.g., amikacin, tigecycline, imipenem, and cefoxitin), administered over 3-12 weeks. This is followed by a continuation phase with a daily oral macrolide (preferably azithromycin), inhaled amikacin, and 2-3 additional oral antibiotics (e.g., moxifloxacin, minocycline, clofazimine, and linezolid) [12–14]. However, MABSC-PD remains a significant therapeutic challenge due to the limited number of safe and effective antibiotics available for treatment, as reflected in poor clinical outcomes with sputum culture conversion rates of approximately 45% [15–16]. Multi-drug resistance towards existing agents creates additional challenges. *M. abscessus* is particularly difficult to treat due to a combination of intrinsic, adaptive, and acquired resistance mechanisms, including permeability barriers, highly efficient drug efflux systems, low-affinity antibiotic targets, and the production of drug-neutralizing enzymes [17–18]. A major driver of resistance is the intrinsic expression of *Bla*_mab_, a broad-spectrum β-lactamase that significantly reduces the activity of β-lactam agents [19]. Recent studies have reported high overall resistance rates to imipenem (55.6%) for MABSC [20]. Notably, imipenem’s efficacy is significantly enhanced when combined with relebactam, a potent inhibitor of *Bla*_mab_, resulting in at least a twofold increase in activity [21–22]. The safety and tolerability of current therapies further complicate treatment efforts. A recent study reported that 79% of patients receiving treatment for MABSC-PD reported adverse side effects, the most severe being ototoxicity, gastrointestinal distress, and myelosuppression. Such side effects – particularly those caused by amikacin, tigecycline, and linezolid, respectively – required therapy modifications for 25% of patients [16]. Thus, there is an urgent need for new therapeutic strategies, including novel antibiotic combinations, to effectively combat MABSC-PD.

Imipenem/cilastatin/relebactam is a novel β-lactam-β-lactamase inhibitor combination that is currently approved for use in adults with hospital-acquired and/or ventilator-associated bacterial pneumonia, complicated urinary tract infections, and complicated intra-abdominal infections. This combination is generally well tolerated, with a relatively low incidence of adverse side effects [23]. Importantly, imipenem/relebactam demonstrates activity against resistant *M. abscessus* strains [21,24–25], with pharmacokinetic data showing effective pulmonary penetration in healthy volunteers [26–27]. Furthermore, imipenem has demonstrated *in vitro* synergistic activity against *M. abscessus* with various antibiotics [21,28–31], and its use in multidrug regimens has been strongly associated with improved treatment outcomes for MABSC-PD, with an adjusted odds ratio of 2.1-2.7 [15–16].

The following study aims to assess the activity of imipenem/relebactam, alongside a selection of antibiotics, against clinical MABSC isolates from PwCF. Given the prevalence of multidrug-resistant MABSC strains in clinical practice, it is critical to assess the efficacy of experimental therapies against contemporary lung disease isolates. Susceptibility testing was conducted on twenty-eight unique MABSC isolates to compare the effectiveness of imipenem/relebactam with ten additional agents. Additionally, checkerboard synergy and time-kill assays were performed to identify optimal antibiotic combinations containing imipenem/relebactam. This research represents the first comprehensive evaluation of imipenem/relebactam in combination with various double and triple antibiotic regimens against CF clinical MABSC isolates, offering valuable insights into potential treatment strategies for MABSC in PwCF.

## 2. Methods

### 2.1. Preparation of antibiotics and growth media

Imipenem/relebactam was provided by Merck. Amikacin, amoxicillin, cefoxitin, cefdinir, cefuroxime, moxifloxacin, azithromycin, rifabutin, clofazimine, minocycline, and tigecycline were purchased from Sigma-Aldrich. Tedizolid was purchased from MedChem Express. Antibiotics were prepared according to their solubility and the manufacturers’ guidelines.

The standard liquid growth medium used for initiating bacterial cultures consisted of Middlebrook 7H9 broth, supplemented with 10% (v/v) Middlebrook oleic acid-albumin-dextrose-catalase (OADC) Enrichment, 0.5% (v/v) glycerol, and 0.05% (v/v) Tween 80. The standard solid growth medium was Middlebrook 7H10 agar enriched with 10% (v/v) OADC. For cryogenic storage, glycerol-stock medium was formulated using Middlebrook 7H9 broth supplemented with 10% (v/v) OADC, 15% (v/v) glycerol, and 0.05% (v/v) Tween 80. Middlebrook 7H9 Broth Base and Middlebrook OADC Enrichment were purchased from Becton, Dickinson, and Company, while Middlebrook 7H10 Agar Base, glycerol, and Tween 80 were purchased from Sigma-Aldrich.

### 2.1. Bacterial strains and culture conditions

The *M. abscessus* reference strain (ATCC 19977) was obtained from the American Type Culture Collection, while CF clinical isolates were acquired from National Jewish Health. Twenty-eight clinical isolates were chosen from the National Jewish Health culture collection to represent the distribution of *M. abscessus* subspecies in PwCF, with selection criteria informed by population genomics data from the Colorado Research and Development Program [32]. For long-term storage, all strains were stored at -80°C in glycerol-stock medium. Frozen stocks were streaked onto round 7H10 plates (100×15 mm polystyrene petri dishes) and incubated at 30°C for three days to promote growth, after which the plates were stored at 2-4°C. Single colonies were then used to inoculate 5 mL cultures of 7H9 broth in 15 mL polypropylene culture tubes. Cultures were incubated at 30°C in a shaking incubator (180 rpm) for 48-96 hours to reach log-phase exponential growth.

### 2.3. Broth microdilution assay and minimum inhibitory concentration (MIC) determination for susceptibility testing

Antibiotic susceptibility testing was performed on the *M. abscessus* ATCC 19977 reference strain and the twenty-eight CF clinical isolates. The antimicrobial agents tested included imipenem/relebactam, imipenem, amikacin, amoxicillin, cefoxitin, cefdinir, cefuroxime, moxifloxacin, azithromycin, tedizolid, rifabutin, clofazimine, minocycline, and tigecycline. The MIC of each antibiotic was determined using the broth microdilution method, according to the Clinical Laboratory Standards Institute (CLSI) guidelines for NTM [33]. Imipenem/relebactam was prepared by serially diluting imipenem while maintaining a fixed concentration of 4 µg/mL relebactam, consistent with prior published studies [24]. Each antibiotic concentration was tested in triplicate. In this study, MIC_50_ was defined as the lowest concentration of antibiotic that inhibited 50% of the tested CF isolates, and MIC_90_ as the lowest concentration of antibiotic that inhibited 90% of the tested CF isolates.

### 2.4. Checkerboard assay and fractional inhibitory concentration (FIC) index determination for synergy testing

Checkerboard assays were conducted to evaluate *in vitro* synergy with imipenem/relebactam against *M. abscessus* ATCC 19977 and the twenty-eight CF clinical isolates, as previously described [30]. The FIC indices were calculated using the following formula: FIC = (MIC_AB_/MIC_A_) + (MIC_BA_/MIC_B_). In this equation, MIC_AB_ is the MIC of drug A tested in combination and MIC_A_ is the MIC of drug A tested alone, while MIC_BA_ is the MIC of drug B tested in combination and MIC_B_ is the MIC of drug B tested alone [34]. An FIC index of ≤ 0.5 indicates synergy, values between > 0.5 to 1 suggest additive effects, values between > 1 to 4 indicate indifference, and values ≥ 4 signify antagonism [34].

Initial screening of antibiotic combinations with imipenem/relebactam for *in vitro* synergy testing against ATCC 19977 included the following agents: amikacin, amoxicillin, cefoxitin, cefdinir, cefuroxime, moxifloxacin, azithromycin, tedizolid, rifabutin, clofazimine, minocycline, and tigecycline. Based on the screening results and a comprehensive literature review, antibiotics demonstrating strong synergy with imipenem/relebactam were prioritized for further testing in clinical isolates. These included cefoxitin, cefdinir, cefuroxime, moxifloxacin, rifabutin, and minocycline. Additionally, amoxicillin, azithromycin, and clofazimine were selected based on their previously reported synergistic potential [21,35–37]. Amikacin and tigecycline were excluded due to significant side effects [16] and the preference for oral regimens, while tedizolid was omitted due to its lack of initial synergy with imipenem/relebactam.

### 2.5. Time-kill assays

To evaluate the overall bactericidal activity of individual antibiotics and synergistic three-drug antibiotic combinations with imipenem/relebactam against *M. abscessus*, time-kill assays were conducted, as previously described [38]. Selection of antibiotics was prioritized based on demonstrated synergy with imipenem/relebactam, activity at clinically achievable concentrations, and alignment with established MABSC treatment guidelines, including cefoxitin, moxifloxacin, azithromycin, rifabutin, and clofazimine. Four concentrations of each antibiotic were selected based on the MICs determined for the *M. abscessus* ATCC 19977 reference strain: 16× MIC, 4× MIC, 1× MIC, and 1/4× MIC. Each antibiotic concentration was tested in duplicate.

Bactericidal activity was defined as a reduction of ≥3 log_10_ CFU/mL compared to the initial count [39]. Synergy was characterized by a ≥2 log_10_ CFU/mL reduction in bacterial load for a given combination compared to the most active single agent. Additivity was defined as a 1 to 2 log_10_ CFU/mL reduction in the final colony count relative to the most active single agent. Indifference was indicated by a change of <1 log_10_ CFU/mL in the final colony count when compared to the most active single agent [39–40].

### 2.6. Data and statistical analysis

Statistical and graphical analyses were performed using GraphPad Prism version 10.1 (GraphPad Software, Inc.). Descriptive analysis of MICs was initially conducted after log_2_-transformation and subsequent correction. The Wilcoxon Signed-Rank test was then applied to log_2_-transformed MICs to compare susceptibility to imipenem alone and imipenem/relebactam. Reductions in bacterial load (log_10_ CFU/mL) were evaluated using one-way ANOVA followed by Tukey’s multiple comparisons test.

## 3. Results

### 3.1. *In Vitro* Susceptibility and Synergy Testing of Antibiotics Against *M. abscessus* ATCC 19977

Susceptibility studies were conducted to evaluate the *in vitro* activities of imipenem/relebactam, as well as amikacin, amoxicillin, cefoxitin, cefdinir, cefuroxime, moxifloxacin, azithromycin, tedizolid, rifabutin, clofazimine, minocycline, and tigecycline against *M. abscessus* ATCC 19977, with results summarized in Table 1.

**Table 1:**
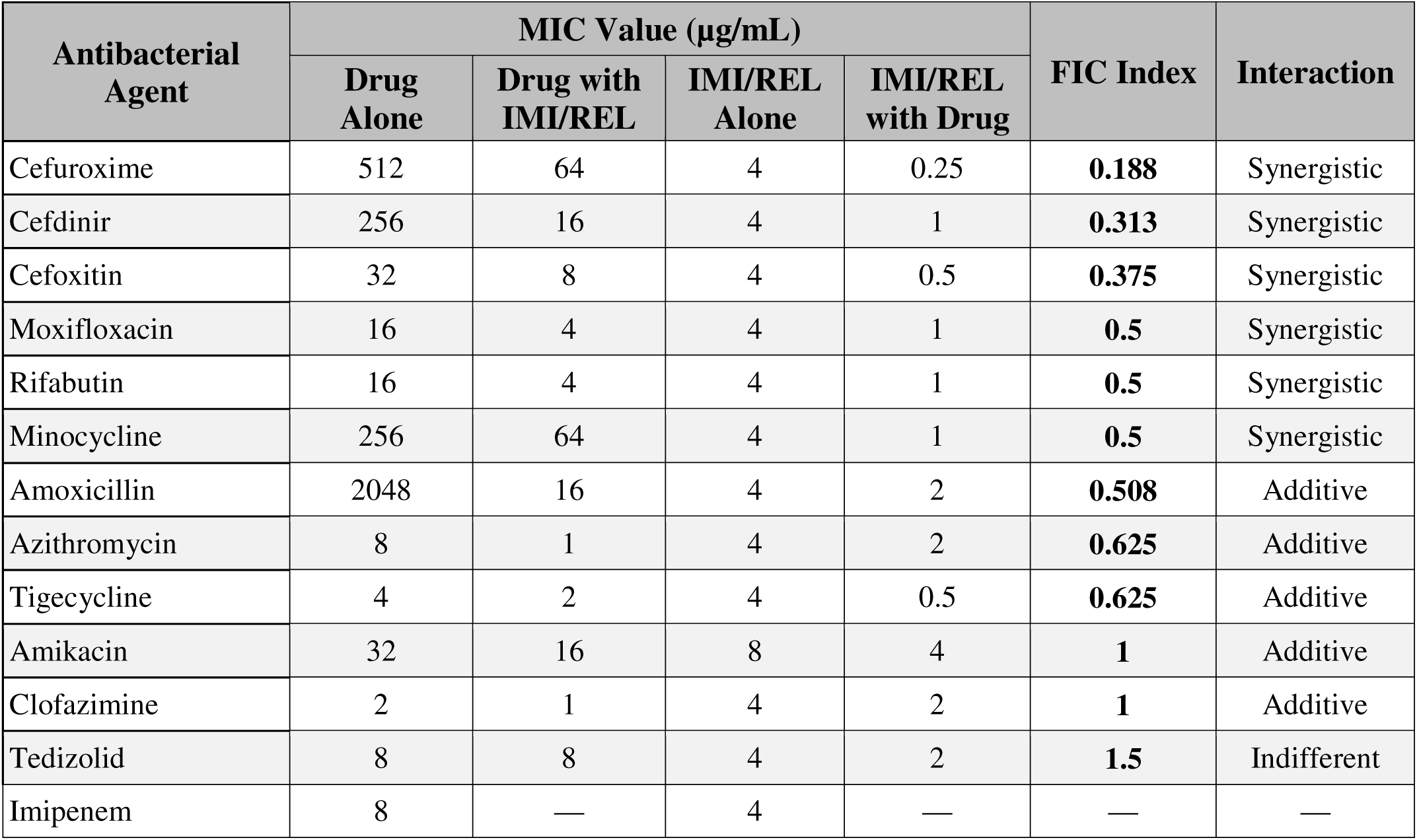
Minimum inhibitory concentrations (MICs) of antibiotics and imipenem/relebactam (IMI/REL), both individually and in combination against the *M. abscessus* ATCC 19977 strain, along with the corresponding susceptibility interpretations, fractional inhibitory concentration (FIC) index values, and their synergism.

| Antibacterial Agent | MIC Value (µg/mL) |  |  |  | FIC Index | Interaction |
| --- | --- | --- | --- | --- | --- | --- |
|  | Drug Alone | Drug with IMI/REL | IMI/REL Alone | IMI/REL with Drug |  |  |
| Cefuroxime | 512 | 64 | 4 | 0.25 | **0.188** | Synergistic |
| Cefdinir | 256 | 16 | 4 | 1 | **0.313** | Synergistic |
| Cefoxitin | 32 | 8 | 4 | 0.5 | **0.375** | Synergistic |
| Moxifloxacin | 16 | 4 | 4 | 1 | **0.5** | Synergistic |
| Rifabutin | 16 | 4 | 4 | 1 | **0.5** | Synergistic |
| Minocycline | 256 | 64 | 4 | 1 | **0.5** | Synergistic |
| Amoxicillin | 2048 | 16 | 4 | 2 | **0.508** | Additive |
| Azithromycin | 8 | 1 | 4 | 2 | **0.625** | Additive |
| Tigecycline | 4 | 2 | 4 | 0.5 | **0.625** | Additive |
| Amikacin | 32 | 16 | 8 | 4 | **1** | Additive |
| Clofazimine | 2 | 1 | 4 | 2 | **1** | Additive |
| Tedizolid | 8 | 8 | 4 | 2 | **1.5** | Indifferent |
| Imipenem | 8 | — | 4 | — | — | — |

Checkerboard assays were conducted to evaluate the *in vitro* synergy of imipenem/relebactam with various antibiotics and FIC indices were calculated to characterize their interactions. The MICs of the antibiotics alone and in combination with imipenem/relebactam against *M. abscessus* ATCC 19977 are also presented in Table 1, along with their respective FIC indices. Synergism with imipenem/relebactam was observed with cefuroxime, cefdinir, cefoxitin, moxifloxacin, rifabutin, and minocycline, while additive effects were noted for amoxicillin, azithromycin, tigecycline, amikacin, and clofazimine. Indifference with imipenem/relebactam was observed with tedizolid, and no antagonism was detected in any of the combinations tested.

### 3.2. *In Vitro* Susceptibility Testing and Screening of Antibiotics for Synergy with Imipenem/Relebactam Against *M. abscessus* CF Clinical Isolates

A summary of the identified *M. abscessus* CF clinical isolates, detailing their subspecies classification and morphology, is provided in Supplementary Table 1. The twenty-eight isolates included 17 (60.7%) *M. abscessus* subsp. *abscessus*, 7 (25.0%) *M. abscessus* subsp. *massiliense*, and 4 (14.3%) *M. abscessus* subsp. *bolletii*. Among these, 15 isolates (53.6%) exhibited the rough morphotype, 7 (25.0%) exhibited the smooth morphotype, and 6 (21.4%) presented intermediate features of both morphotypes, leading to their classification as “unknown” along the smooth-rough spectrum.

Susceptibility and synergy testing were performed to evaluate the *in vitro* activity of imipenem/relebactam in combination with various antibiotics against these isolates. The MIC_50_ values for the individual antibiotics, both alone and in combination with imipenem/relebactam, along with the median FIC indices, are summarized in Table 2. Supplementary Tables 2-10 provide these MIC values and FIC indices for each individual CF clinical isolate strain, along with the MIC_50_ and MIC_90_ values.

**Table 2:** Susceptibility and synergy of antibiotics with imipenem/relebactam (IMI/REL) against *M. abscessus* CF clinical isolates (*n*=28)

| Antibacterial Agent | MIC<sub>50</sub> Value (µg/mL)<sup>a</sup> |  |  |  | FIC Index | Interaction |
| --- | --- | --- | --- | --- | --- | --- |
|  | Drug Alone | Drug with IMI/REL | IMI/REL Alone | IMI/REL with Drug |  |  |
| Cefuroxime | 512<br>(256, 512) | 64<br>(32, 128) | 4<br>(4, 8) | 0.38<br>(0.13, 0.50) | **0.250**<br>**(0.180, 0.328)** | Synergistic |
| Cefdinir | 192<br>(112, 256) | 24<br>(14, 40) | 4<br>(4, 8) | 1<br>(0.50, 1) | **0.375**<br>**(0.313, 0.502)** | Synergistic |
| Amoxicillin | 2048<br>(1024, 2048) | 128<br>(32, 256) | 8<br>(4, 8) | 2<br>(1, 2.50) | **0.438**<br>**(0.352, 0.625)** | Synergistic |
| Cefoxitin | 48<br>(32, 64) | 8<br>(4, 8) | 8<br>(4, 16) | 2<br>(1, 8) | **0.500**<br>**(0.375, 0.563)** | Synergistic |
| Rifabutin | 8<br>(7, 16) | 2<br>(1, 4) | 4<br>(4, 8) | 1<br>(0.88, 2) | **0.563**<br>**(0.500, 0.625)** | Additive |
| Azithromycin | 8<br>(4, 16) | 2<br>(1, 4) | 8<br>(8, 16) | 2<br>(1, 4) | **0.625**<br>**(0.500, 0.750)** | Additive |
| Moxifloxacin | 16<br>(8, 16) | 4<br>(4, 4) | 6<br>(4, 8) | 2<br>(2, 2) | **0.750**<br>**(0.500, 0.750)** | Additive |
| Clofazimine | 1<br>(0.88, 2) | 0.50<br>(0.25, 0.63) | 8<br>(4, 8) | 2<br>(1, 4) | **0.750**<br>**(0.625, 1.000)** | Additive |
| Minocycline | 256<br>(256, 256) | 96<br>(64, 128) | 4<br>(2, 5) | 2<br>(1, 2) | **0.750**<br>**(0.609, 1.039)** | Additive |

<sup>a</sup> Data are presented as MIC<sub>50</sub> (interquartile range [IQR]).

The antibiotic combinations with imipenem/relebactam demonstrated varying degrees of synergistic, additive, and indifferent effects across the tested CF clinical isolates (Figure 1). However, the median MIC and FIC values for the CF clinical isolates were closely aligned with the corresponding values from the ATCC 19977 strain, indicating similar overall levels of susceptibility and synergy to the antibiotics tested (Tables 1 and 2). Specifically, median FIC indices revealed that, in the clinical isolates, imipenem/relebactam exhibited synergy with cefuroxime, cefdinir, cefoxitin, and amoxicillin, and additive effects with rifabutin, azithromycin, moxifloxacin, clofazimine, and minocycline.

**Figure 1:**
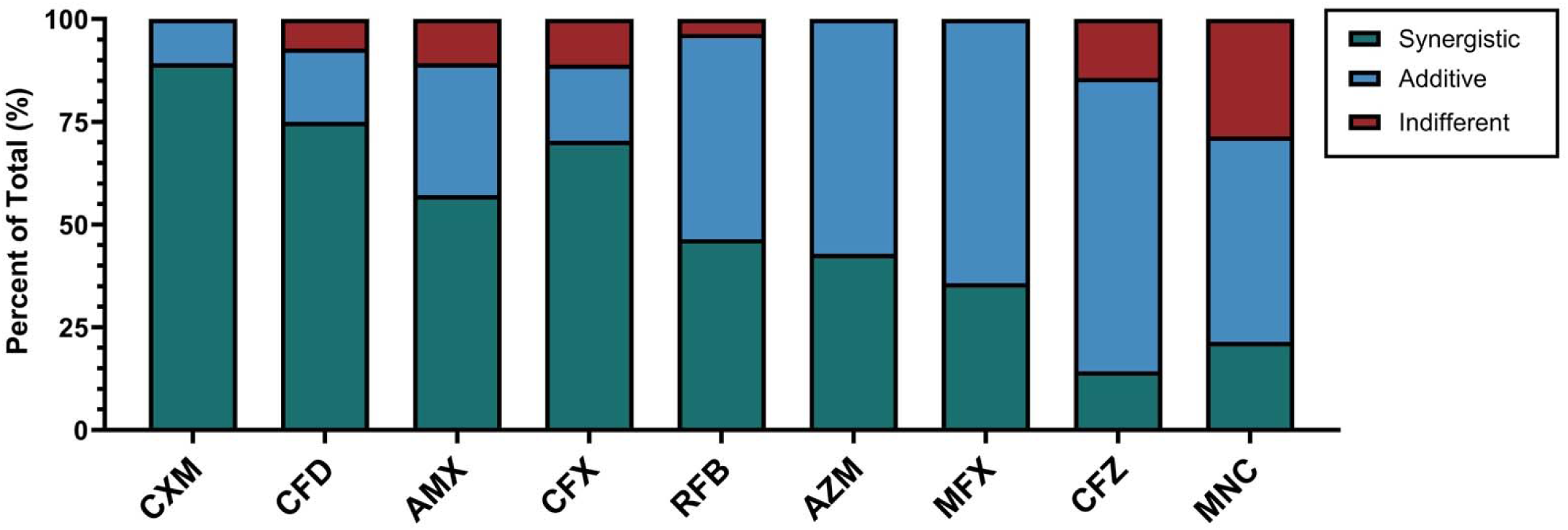
Percentages of synergistic, additive, and indifferent effects of antibiotics combined with imipenem/relebactam in *M. abscessus* CF clinical isolates CXM = cefuroxime, CFD = cefdinir, AMX = amoxicillin, CFX = cefoxitin, RFB = rifabutin, AZM = azithromycin, MFX = moxifloxacin, CFZ = clofazimine, MNC = minocycline

Additionally, comparisons between imipenem alone and imipenem/relebactam were conducted to assess the effect of the β-lactamase inhibitor on imipenem’s activity against *M. abscessus* (Supplementary Table 11). A significant difference in MIC values was observed, with imipenem alone exhibiting higher MICs (*p* = 0.0054). The MIC_50_ was 8 µg/mL for both treatments, while the MIC_90_ values were 9.2 µg/mL for imipenem/relebactam and 16.0 µg/mL for imipenem alone.

### 3.3. Evaluation of Three-Drug Antibiotic Combinations with Imipenem/Relebactam Using Time-Kill Assay

3.3.1. Initial Screening of Antibiotic Combinations Against *M. abscessus* ATCC 19977

An initial endpoint activity assay was performed to assess the efficacy of six individual antibiotics and seven combination treatments with imipenem/relebactam, all administered at 1× MIC. The mean bacterial loads (log_10_ CFU/mL) over 72 hours for these single and combination therapies are presented in Figure 2*A-B*. None of the individual antibiotics tested (imipenem/relebactam, cefoxitin, moxifloxacin, azithromycin, rifabutin, clofazimine) exhibited bactericidal activity at 72 hours, but all showed significant reductions in bacterial load compared to the control, with clofazimine being the most effective (diff = −2.90, p < 0.0001) (Fig. 2*A*). Several combinations with imipenem/relebactam demonstrated bactericidal activity and significant bacterial reductions, including combinations with cefoxitin and rifabutin (diff = −6.43, p < 0.0001), cefoxitin and moxifloxacin (diff = −4.26, p < 0.0001), and azithromycin and clofazimine (diff = −3.89, *p* = 0.0011) (Fig. 2*B*).

**Figure 2:**
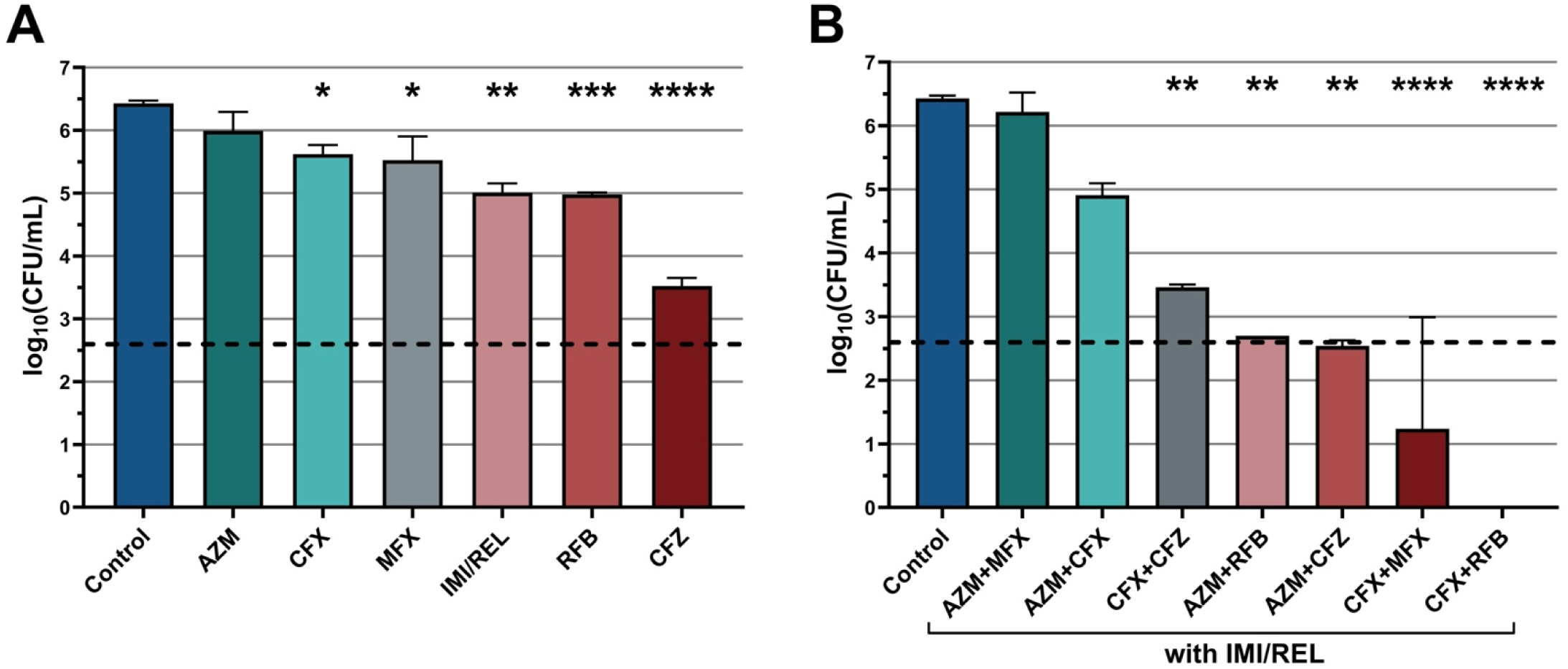
Mean bacterial loads (log_10_ CFU/mL) of *M. abscessus* ATCC 1997 over 72 hours with (A) single-agent therapies and (B) three-drug combination therapies with imipenem/relebactam (IMI/REL) Data are presented as mean with standard errors of the mean. Asterisks denote significant differences compared to the untreated control. The horizontal dashed line marks a 3-log reduction in CFU/mL relative to the initial count of the untreated control, denoting bactericidal activity. (*p<0.05, **p<0.01, ***p<0.001, ****p<0.0001, post-hoc Tukey’s HSD test) AZM = azithromycin, CFX = cefoxitin, MFX = moxifloxacin, RFB = rifabutin, CFZ = clofazimine

#### 3.3.2. Kinetic Time-Kill Assay with Antibiotic Combinations Against *M. abscessus* ATCC 19977 and CF Clinical Isolates CF13 and CF258

Kinetic time-kill assays were conducted to evaluate the efficacy of four imipenem/relebactam combination treatments, selected for their bactericidal activity in initial screening assays. Both the combinations and individual antibiotics were tested against the *M. abscessus* ATCC 19977 reference strain at 1× MIC, with bacterial loads (log_10_ CFU/mL) at 24, 48, and 72 hours shown in Figure 3. All treatments significantly reduced bacterial load compared to the control at 72 hours. Imipenem/relebactam alone slightly reduced bacterial load compared to the initial inoculum, although this difference was not statistically significant. Combination therapies exhibited enhanced efficacy. Imipenem/relebactam combinations with cefoxitin and rifabutin, as well as with cefoxitin and moxifloxacin, achieved bactericidal and synergistic effects, resulting in complete eradication. Other combinations were bacteriostatic. Similar trends were observed in assays at 16× MIC, 4× MIC, and 1/4× MIC (Supplementary Fig. 1).

**Figure 3:**
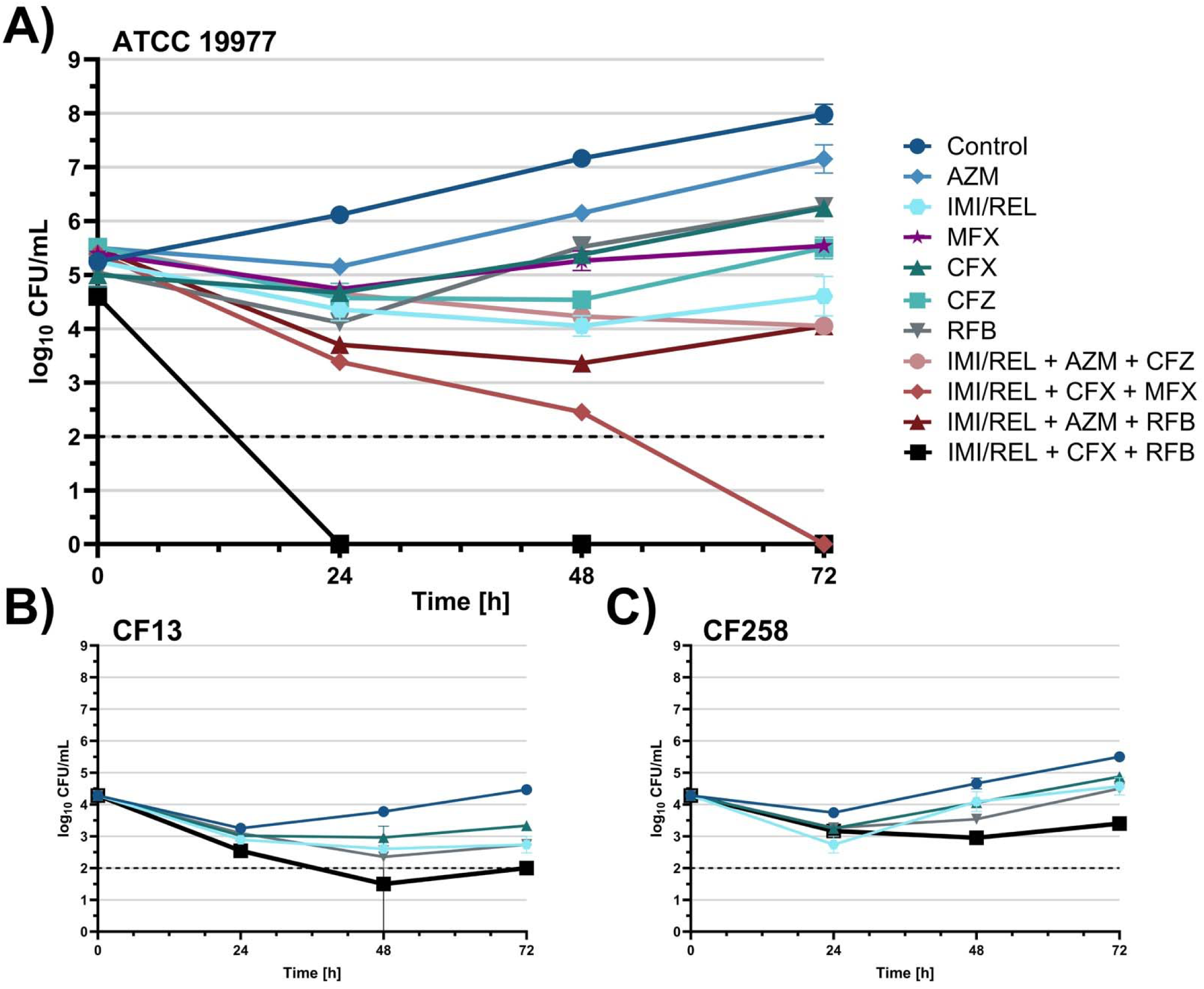
A) Bacterial load (log_10_ CFU/ml) of *M. abscessu*s ATCC 19977 over 72 hours with single-agent therapies and three-drug combination therapies with imipenem/relebactam (IMI/REL) at 1× MIC, B) *M. abscessus* CF isolate 13 bacterial load over 72 hours with IMI/REL, cefoxitin, rifabutin, and their combination at 1× MIC. C) *M. abscessus* CF isolate 258 bacterial load over 72 hours with the same treatments as in B. Data are presented as mean with standard errors of the mean. The horizontal dashed line marks the lower limit of detection. AZM = azithromycin, MFX = moxifloxacin, CFX = cefoxitin, CFZ = clofazimine, RFB = rifabutin

Clinical isolates CF13 and CF258 were selected for additional kinetic time-kill assays with imipenem/relebactam, cefoxitin, and rifabutin based on their susceptibility to imipenem, representing MIC_50_ (moderately susceptible) and MIC_90_ (least susceptible) strains, respectively (Figure 3*B-C*, Supplementary Figs. 2 and 3). Across all concentrations, the imipenem/relebactam, cefoxitin, and rifabutin combination achieved the greatest bacterial reduction in both isolates. In CF13, all treatments were bactericidal at 16× MIC, with rifabutin and the combination also achieving eradication at 4× MIC. Although none were bactericidal at 1× MIC, all significantly reduced bacterial load compared to the initial inoculum. At 1/4× MIC, rifabutin (*p* = 0.0093) and the combination (*p* = 0.0045) retained this activity. In CF258, cefoxitin, rifabutin, and three-drug combination therapy were bactericidal, with rifabutin and the combination achieving eradication at 16× MIC. No treatments were bactericidal at 4×, 1×, or 1/4× MIC, but all significantly reduced bacterial load at 4× MIC compared to the initial inoculum. At 1× MIC, only the imipenem/relebactam, cefoxitin, and rifabutin combination significantly reduced bacterial load from the initial count (*p* = 0.0100).

## 4. Discussion

This study explores the therapeutic potential of imipenem/relebactam in combination with various antibiotics for treating *M. abscessus*, a key pathogen responsible for severe lung infections and adverse clinical outcomes in PwCF. Utilizing a diverse array of clinical isolates from CF patients alongside a reference strain, we evaluated the *in vitro* efficacy of antibiotic combinations containing imipenem/relebactam through susceptibility and synergy testing. Our findings were further substantiated by time-kill kinetic assays. These results reveal substantial synergistic interactions between imipenem/relebactam and antibiotics from multiple drug classes, indicating *in vivo* studies are warranted to evaluate clinical outcomes using these combinations for the treatment of these challenging infections.

Our results confirm that imipenem/relebactam exhibits superior *in vitro* efficacy over imipenem alone against *M. abscessus* ATCC 19977 and CF clinical isolates. This enhanced activity can be attributed to relebactam, a β-lactamase inhibitor that enhances the stability and activity of imipenem against β-lactamase-producing *M. abscessus* strains [19,22,25]. The combination of imipenem/relebactam significantly reduced MIC values compared to imipenem monotherapy, with a fold change of approximately 1.74 in the MIC_90_ values (9.2 µg/mL vs 16.0 µg/mL, respectively), reinforcing its potential as a promising treatment for *M. abscessus* infections.

Susceptibility testing conducted on the *M. abscessus* ATCC 19977 reference strain showed MIC values consistent with those reported in the literature for imipenem, amoxicillin, moxifloxacin, minocycline, amikacin, cefoxitin, cefdinir, cefuroxime, azithromycin, tedizolid, and rifabutin [24,41–45]. However, greater variability was observed for clofazimine, with MIC values 4 to 8-fold higher, and for tigecycline, which showed MIC values 8 to 32-fold higher than published studies [42,46–47]. Factors potentially contributing to MIC variability include differences in inoculum calibration, drug stability, inherent antibiotic variability, variations in laboratory techniques, environmental conditions during testing, and potential genetic drift within bacterial populations across studies. To mitigate these challenges, all susceptibility testing for *M. abscessus* ATCC 19977 and CF clinical isolates was conducted following CLSI guidelines, ensuring experimental consistency and facilitating more accurate comparisons with literature values [33].

Standard susceptibility testing was also performed on CF clinical isolates. Consistent with previous studies highlighting the genotypic and phenotypic diversity of *M. abscessus* [48–49], we observed substantial heterogeneity in MIC values and antibiotic responses among the CF isolates. This variability may stem from the genetic diversity of *M. abscessus* strains within the CF population, as well as bacterial adaptations to the CF lung environment [50–52]. These findings underscore the importance of considering this heterogeneity in the development of targeted treatment strategies for *M. abscessus* infections in PwCF.

Checkerboard analyses of CF clinical isolates revealed synergistic interactions between imipenem/relebactam and cefuroxime, cefdinir, amoxicillin, and cefoxitin, aligning with established findings on β-lactam synergy [30–31,53]. Rifabutin, azithromycin, moxifloxacin, clofazimine, and minocycline also demonstrated additive effects with imipenem/relebactam. However, cefuroxime, cefdinir, amoxicillin, and minocycline are unlikely to achieve clinically relevant plasma or intracellular concentrations under standard dosing regimens [54–57], limiting their potential efficacy alone or in combination with imipenem/relebactam against *M. abscessus*. Time-kill assays further supported our findings, showing that the greatest reductions in bacterial load occurred when imipenem/relebactam was combined with cefoxitin and moxifloxacin or rifabutin.

These results highlight the *in vitro* efficacy of multi-drug regimens—particularly those incorporating imipenem/relebactam with cefoxitin, azithromycin, moxifloxacin, clofazimine, and rifabutin—for effectively targeting *M. abscessus* infections. Cefoxitin, a guideline-recommended intravenous antibiotic for the intensive phase of *M. abscessus* therapy [13], exhibited robust efficacy and consistent synergy with imipenem/relebactam, in line with previous findings that reinforce its ongoing relevance in *M. abscessus* management [58]. Azithromycin, the preferred oral macrolide for both intensive and continuation phases [13], is known for its anti-inflammatory and immunomodulatory properties in PwCF [59–60], and its use in multidrug regimens has been associated with improved outcomes in MABSC-PD treatment [15].

Moxifloxacin, recommended during the continuation phase and available orally [13], has also demonstrated promising preclinical anti-inflammatory activity in CF bronchial cell models [61]. Among emerging agents, clofazimine, a key drug used in the treatment of leprosy, has been increasingly used for nontuberculosis mycobacterial infections, particularly those caused by *M. abscessus* [62–63]. Rifabutin remains the only rifamycin shown to be effective against *M. abscessus* [45,64–65], and its inclusion in multi-drug regimens is further supported by its ability to reduce macrolide resistance [66].

While this study provides valuable insights, there are several limitations to consider. First, the treatment history of tested isolates was unknown and previous exposure to antimicrobials (e.g., macrolides) may have affected drug susceptibility testing results [67]. Furthermore, not all antibiotics available for *M. abscessus* treatment were evaluated in this study. While imipenem/relebactam was tested alongside twelve different antibiotics, agents such as linezolid and bedaquiline were omitted due to a lack of documented synergy with other agents. The stability of certain compounds, particularly imipenem/relebactam, presents a potential concern. Certain β-lactams are known to be relatively unstable, with one study reporting a degradation half-life of 16.9 hours for imipenem in media [68]. To evaluate the potential impact of this instability on treatment efficacy, we tested imipenem/relebactam monotherapy with and without media replacement every 24 hours. Media replacement with imipenem/relebactam resulted in a roughly 1 log_10_ greater reduction in bacterial load (CFU/mL) compared to the non-replacement group at 16×, 4×, and 1× MIC concentrations, although no statistically significant differences were observed between the groups. Last, several previous studies have highlighted that the *in vitro* efficacy of antibiotics against *M. abscessus* may not consistently translate to *in vivo* outcomes [69]. The static concentrations used in this study (16×, 4×, 1×, and 1/4× MIC) are typical for *in vitro* testing, but may not fully reflect dynamic pharmacokinetics in a clinical setting. Future studies should utilize dynamic *in vitro* models [70] and *in vivo* animal models [71–72] to better simulate human conditions and confirm the clinical relevance of these findings. Despite these limitations, the large-scale use of clinical isolates and antibiotic combinations in this study provides valuable data that may otherwise require substantial time and resources in more complex models.

## 5. Conclusion

In conclusion, imipenem/relebactam demonstrated potent *in vitro* activity against clinical isolates of MABSC, with substantial synergistic interactions observed when combined with antibiotics from multiple drug classes. These findings suggest that imipenem/relebactam may offer a significant advancement in the management of infections involving MABSC in PwCF. Moreover, the promising efficacy of multidrug regimens pairing imipenem/relebactam with agents such as cefoxitin, azithromycin, moxifloxacin, clofazimine, and rifabutin highlights potential therapeutic strategies for targeting *M. abscessus*. Future preclinical studies are warranted to validate these findings, optimize drug combinations, and evaluate their effectiveness in treating MABSC pulmonary disease.

## Supporting information

Supplementary Electronic Materials

## 7. Acknowledgments

We would like to acknowledge National Jewish Health for providing the *M. abscessus* CF clinical isolates used in this study. This study was supported by an investigator-initiated grant from Merck & Co., Inc, and by NIH/NIDCR training grant T90-DE02198213.

