## Supplementary Electronic Materials for "*In vitro* activity of imipenem/relebactam alone and in combination against cystic fibrosis isolates of *Mycobacterium abscessus*"

**9. Supplementary:**

**Supplementary Table 1**: *M. abscessus* CF clinical isolates identified according to their subspecies classification and morphology

| **CF Patient ASID** | **WGS**  **Identification** | **CF Morphology** |
| --- | --- | --- |
| CF00006 | MAB | R |
| CF00013 | MAB | R |
| CF00016 | MAB | R |
| CF00017 | MAB | R |
| CF00023 | MAB | R |
| CF00038 | MAB | S |
| CF00040 | MAB | R |
| CF00041 | MAB | S |
| CF00043 | MAB | R |
| CF00136 | MAB | unknown |
| CF00258 | MAB | unknown |
| CF00855 | MAB | unknown |
| CF01975 | MAB | R |
| CF02033 | MAB | R |
| CF02279 | MAB | R |
| CF02319 | MAB | R |
| CF02486 | MAB | S |
| CF00008 | MMAS | R |
| CF00030 | MMAS | S |
| CF00035 | MMAS | R |
| CF00042 | MMAS | S |
| CF00046 | MMAS | unknown |
| CF00047 | MMAS | S |
| CF00883 | MMAS | R |
| CF00020 | MBOL | S |
| CF00113 | MBOL | unknown |
| CF00868 | MBOL | unknown |
| CF02061 | MBOL | R |

MAB, *M. abscessus* subsp. *abscessus*; MBOL, *M. abscessus* subsp. *bolletii*; MMAS, *M. abscessus* subsp. *massiliense*; R, rough; S, smooth

**Supplementary Table 2**: Minimum inhibitory concentrations (MICs) of amoxicillin and imipenem/relebactam (IMI/REL), both individually and in combination against the *M. abscessus* CF clinical isolates, along with the corresponding fractional inhibitory concentration (FIC) index values and their synergism

| **WGS ID** | **CF Patient ID** | **MIC Value (µg/mL)** | | | | **FIC Index** | **Interaction** |
| --- | --- | --- | --- | --- | --- | --- | --- |
|  |  | **Amoxicillin Alone** | **Amoxicillin with IMI/REL** | **IMI/REL**  **Alone** | **IMI/REL with Amoxicillin** |  |  |
| ***M.***  ***abscessus*** | CF00006 | 2048 | 256 | 16 | 8 | **0.625** | Additive |
|  | CF00013 | 2048 | 512 | 2 | 0.5 | **0.500** | Synergistic |
|  | CF00016 | 2048 | 512 | 8 | 2 | **0.500** | Synergistic |
|  | CF00017 | 1024 | 64 | 4 | 2 | **0.563** | Additive |
|  | CF00023 | 2048 | 64 | 4 | 0.5 | **0.156** | Synergistic |
|  | CF00038 | 2048 | 16 | 4 | 1 | **0.258** | Synergistic |
|  | CF00040 | 2048 | 16 | 8 | 8 | **1.008** | Indifferent |
|  | CF00041 | 2048 | 256 | 4 | 1 | **0.375** | Synergistic |
|  | CF00043 | 2048 | 256 | 4 | 1 | **0.375** | Synergistic |
|  | CF00136 | 2048 | 512 | 8 | 1 | **0.375** | Synergistic |
|  | CF00258 | 2048 | 256 | 8 | 4 | **0.625** | Additive |
|  | CF00855 | 2048 | 256 | 4 | 1 | **0.375** | Synergistic |
|  | CF01975 | 1024 | 128 | 16 | 4 | **0.375** | Synergistic |
|  | CF02033 | 2048 | 256 | 8 | 2 | **0.375** | Synergistic |
|  | CF02279 | 2048 | 64 | 4 | 2 | **0.531** | Additive |
|  | CF02319 | 2048 | 8 | 2 | 2 | **1.004** | Indifferent |
|  | CF02486 | 1024 | 256 | 8 | 4 | **0.750** | Additive |
| ***M. massiliense*** | CF00008 | 2048 | 8 | 4 | 2 | **0.504** | Additive |
|  | CF00030 | 2048 | 64 | 8 | 0.5 | **0.094** | Synergistic |
|  | CF00035 | 2048 | 2048 | 8 | 1 | **1.125** | Indifferent |
|  | CF00042 | 1024 | 32 | 16 | 4 | **0.281** | Synergistic |
|  | CF00046 | 1024 | 256 | 16 | 8 | **0.750** | Additive |
|  | CF00047 | 2048 | 32 | 8 | 2 | **0.266** | Synergistic |
|  | CF00883 | 512 | 16 | 8 | 0.125 | **0.047** | Synergistic |
| ***M.***  ***bolletii*** | CF00020 | 1024 | 512 | 8 | 2 | **0.750** | Additive |
|  | CF00113 | 1024 | 128 | 8 | 2 | **0.375** | Synergistic |
|  | CF00868 | 2048 | 64 | 8 | 2 | **0.281** | Synergistic |
|  | CF02061 | 2048 | 32 | 4 | 2 | **0.516** | Additive |
| **MIC_50_** | | 2048 | 128 | 8 | 2 | — | — |
| **MIC_90_** | | 2048 | 512 | 16 | 4.9 | — | — |

**Supplementary Table 3**: Minimum inhibitory concentrations (MICs) of cefoxitin and imipenem/relebactam (IMI/REL), both individually and in combination against the *M. abscessus* CF clinical isolates, along with the corresponding fractional inhibitory concentration (FIC) index values and their synergism

| **WGS ID** | **CF Patient ID** | **MIC Value (µg/mL)** | | | | **FIC Index** | **Interaction** |
| --- | --- | --- | --- | --- | --- | --- | --- |
|  |  | **Cefoxitin Alone** | **Cefoxitin with IMI/REL** | **IMI/REL Alone** | **IMI/REL with Cefoxitin** |  |  |
| ***M. abscessus*** | CF00006 | 64 | 4 | 16 | 16 | **1.063** | Indifferent |
|  | CF00013 | 128 | 32 | 8 | 2 | **0.500** | Synergistic |
|  | CF00016 | 64 | 16 | 8 | 2 | **0.500** | Synergistic |
|  | CF00017 | 32 | 8 | 4 | 2 | **0.750** | Additive |
|  | CF00023 | 64 | 16 | 4 | 1 | **0.500** | Synergistic |
|  | CF00038 | 32 | 8 | 16 | 2 | **0.375** | Synergistic |
|  | CF00040 | 128 | 8 | 32 | 32 | **1.063** | Indifferent |
|  | CF00041 | 32 | 8 | 4 | 1 | **0.500** | Synergistic |
|  | CF00043 | 64 | 8 | 4 | 1 | **0.375** | Synergistic |
|  | CF00136 | 32 | 8 | 4 | 1 | **0.500** | Synergistic |
|  | CF00258 | 64 | 8 | 8 | 1 | **0.250** | Synergistic |
|  | CF00855 | 128 | 8 | 64 | 32 | **0.563** | Additive |
|  | CF01975 | 32 | 4 | 64 | 8 | **0.250** | Synergistic |
|  | CF02033 | 64 | 8 | 32 | 4 | **0.250** | Synergistic |
|  | CF02279 | 64 | 8 | 4 | 1 | **0.375** | Synergistic |
|  | CF02319 | 64 | 2 | 8 | 2 | **0.281** | Synergistic |
|  | CF02486 | 32 | 4 | 8 | 2 | **0.375** | Synergistic |
| ***M. massiliense*** | CF00008 | 64 | 4 | 16 | 8 | **0.563** | Additive |
|  | CF00030 | 32 | 4 | 16 | 4 | **0.375** | Synergistic |
|  | CF00035 | 32 | 4 | 8 | 8 | **1.125** | Indifferent |
|  | CF00042 | 16 | 4 | 16 | 16 | **1.250** | Indifferent |
|  | CF00046 | 64 | 4 | 16 | 8 | **0.563** | Additive |
|  | CF00047 | 32 | 4 | 4 | 1 | **0.375** | Synergistic |
|  | CF00883 | 16 | 4 | 8 | 4 | **0.750** | Additive |
| ***M.***  ***bolletii*** | CF00020 | 32 | 4 | 16 | 4 | **0.375** | Synergistic |
|  | CF00113 | 64 | 8 | 4 | 1 | **0.375** | Synergistic |
|  | CF00868 | 32 | 8 | 8 | 2 | **0.500** | Synergistic |
|  | CF02061 | 32 | 8 | 4 | 1 | **0.500** | Synergistic |
| **MIC_50_**  2048 | | 45.3 | 8 | 8 | 2 | — | — |
| **MIC_90_**  2048 | | 78.8 | 9.8 | 32 | 16 | — | — |

**Supplementary Table 4**: Minimum inhibitory concentrations (MICs) of cefdinir and imipenem/relebactam (IMI/REL), both individually and in combination against the *M. abscessus* CF clinical isolates, along with the corresponding fractional inhibitory concentration (FIC) index values and their synergism

| **WGS ID** | **CF Patient ID** | **MIC Value (µg/mL)** | | | | **FIC Index** | **Interaction** |
| --- | --- | --- | --- | --- | --- | --- | --- |
|  |  | **Cefdinir**  **Alone** | **Cefdinir with IMI/REL** | **IMI/REL**  **Alone** | **IMI/REL with Cefdinir** |  |  |
| ***M.***  ***abscessus*** | CF00006 | 256 | 128 | 16 | 0.125 | **0.508** | Additive |
|  | CF00013 | 256 | 32 | 4 | 1 | **0.375** | Synergistic |
|  | CF00016 | 128 | 16 | 4 | 1 | **0.375** | Synergistic |
|  | CF00017 | 256 | 8 | 2 | 0.5 | **0.281** | Synergistic |
|  | CF00023 | 256 | 32 | 4 | 1 | **0.375** | Synergistic |
|  | CF00038 | 256 | 64 | 4 | 1 | **0.500** | Synergistic |
|  | CF00040 | 256 | 128 | 8 | 1 | **0.625** | Additive |
|  | CF00041 | 256 | 32 | 8 | 1 | **0.250** | Synergistic |
|  | CF00043 | 128 | 8 | 4 | 0.5 | **0.188** | Synergistic |
|  | CF00136 | 16 | 16 | 16 | 0.5 | **1.031** | Indifferent |
|  | CF00258 | 256 | 128 | 16 | 2 | **0.625** | Additive |
|  | CF00855 | 64 | 16 | 4 | 0.5 | **0.375** | Synergistic |
|  | CF01975 | 256 | 32 | 8 | 0.125 | **0.141** | Synergistic |
|  | CF02033 | 256 | 64 | 4 | 1 | **0.500** | Synergistic |
|  | CF02279 | 512 | 32 | 2 | 0.25 | **0.188** | Synergistic |
|  | CF02319 | 128 | 8 | 4 | 1 | **0.313** | Synergistic |
|  | CF02486 | 256 | 16 | 4 | 1 | **0.313** | Synergistic |
| ***M. massiliense*** | CF00008 | 256 | 64 | 8 | 1 | **0.375** | Synergistic |
|  | CF00030 | 128 | 8 | 4 | 1 | **0.313** | Synergistic |
|  | CF00035 | 128 | 16 | 8 | 1 | **0.250** | Synergistic |
|  | CF00042 | 16 | 16 | 8 | 0.125 | **1.016** | Indifferent |
|  | CF00046 | 256 | 64 | 8 | 0.5 | **0.313** | Synergistic |
|  | CF00047 | 128 | 32 | 4 | 1 | **0.500** | Synergistic |
|  | CF00883 | 8 | 2 | 4 | 2 | **0.750** | Additive |
| ***M.***  ***bolletii*** | CF00020 | 64 | 8 | 4 | 1 | **0.375** | Synergistic |
|  | CF00113 | 64 | 8 | 4 | 2 | **0.625** | Additive |
|  | CF00868 | 128 | 32 | 8 | 1 | **0.375** | Synergistic |
|  | CF02061 | 64 | 16 | 4 | 1 | **0.500** | Synergistic |
| **MIC_50_** | | 181 | 22.6 | 4 | 1 | — | — |
| **MIC_90_** | | 256 | 78.8 | 9.8 | 1.2 | — | — |

**Supplementary Table 5**: Minimum inhibitory concentrations (MICs) of cefuroxime and imipenem/relebactam (IMI/REL), both individually and in combination against the *M. abscessus* CF clinical isolates, along with the corresponding fractional inhibitory concentration (FIC) index values and their synergism

| **WGS ID** | **CF Patient ID** | **MIC Value (µg/mL)** | | | | **FIC Index** | **Interaction** |
| --- | --- | --- | --- | --- | --- | --- | --- |
|  |  | **Cefuroxime**  **Alone** | **Cefuroxime**  **with IMI/REL** | **IMI/REL**  **Alone** | **IMI/REL with Cefuroxime** |  |  |
| ***M.***  ***abscessus*** | CF00006 | 512 | 128 | 8 | 0.5 | **0.313** | Synergistic |
|  | CF00013 | 512 | 128 | 4 | 0.125 | **0.281** | Synergistic |
|  | CF00016 | 256 | 64 | 16 | 0.5 | **0.281** | Synergistic |
|  | CF00017 | 512 | 32 | 4 | 0.5 | **0.188** | Synergistic |
|  | CF00023 | 512 | 64 | 4 | 0.5 | **0.250** | Synergistic |
|  | CF00038 | 64 | 16 | 4 | 0.125 | **0.281** | Synergistic |
|  | CF00040 | 512 | 64 | 8 | 0.5 | **0.188** | Synergistic |
|  | CF00041 | 256 | 64 | 4 | 0.25 | **0.313** | Synergistic |
|  | CF00043 | 256 | 16 | 4 | 0.125 | **0.094** | Synergistic |
|  | CF00136 | 128 | 16 | 16 | 0.125 | **0.133** | Synergistic |
|  | CF00258 | 256 | 64 | 8 | 1 | **0.375** | Synergistic |
|  | CF00855 | 512 | 128 | 8 | 0.5 | **0.313** | Synergistic |
|  | CF01975 | 256 | 128 | 16 | 1 | **0.563** | Additive |
|  | CF02033 | 1024 | 256 | 4 | 0.5 | **0.375** | Synergistic |
|  | CF02279 | 512 | 32 | 2 | 0.25 | **0.188** | Synergistic |
|  | CF02319 | 512 | 64 | 4 | 0.5 | **0.250** | Synergistic |
|  | CF02486 | 512 | 32 | 2 | 0.125 | **0.125** | Synergistic |
| ***M. massiliense*** | CF00008 | 1024 | 128 | 4 | 0.125 | **0.156** | Synergistic |
|  | CF00030 | 1024 | 64 | 4 | 0.125 | **0.094** | Synergistic |
|  | CF00035 | 512 | 64 | 4 | 0.25 | **0.188** | Synergistic |
|  | CF00042 | 64 | 32 | 16 | 0.25 | **0.516** | Additive |
|  | CF00046 | 256 | 32 | 4 | 0.25 | **0.188** | Synergistic |
|  | CF00047 | 256 | 64 | 4 | 0.5 | **0.375** | Synergistic |
|  | CF00883 | 1024 | 512 | 16 | 2 | **0.625** | Additive |
| ***M.***  ***bolletii*** | CF00020 | 512 | 64 | 8 | 0.25 | **0.156** | Synergistic |
|  | CF00113 | 2048 | 256 | 8 | 2 | **0.375** | Synergistic |
|  | CF00868 | 128 | 16 | 4 | 0.125 | **0.156** | Synergistic |
|  | CF02061 | 512 | 64 | 4 | 0.5 | **0.250** | Synergistic |
| **MIC_50_** | | 512 | 64 | 4 | 0.4 | — | — |
| **MIC_90_** | | 1024 | 157.6 | 16 | 1 | — | — |

**Supplementary Table 6**: Minimum inhibitory concentrations (MICs) of moxifloxacin and imipenem/relebactam (IMI/REL), both individually and in combination against the *M. abscessus* CF clinical isolates, along with the corresponding fractional inhibitory concentration (FIC) index values and their synergism

| **WGS ID** | **CF Patient ID** | **MIC Value (µg/mL)** | | | | **FIC Index** | **Interaction** |
| --- | --- | --- | --- | --- | --- | --- | --- |
|  |  | **Moxifloxacin**  **Alone** | **Moxifloxacin**  **with IMI/REL** | **IMI/REL**  **Alone** | **IMI/REL with Moxifloxacin** |  |  |
| ***M.***  ***abscessus*** | CF00006 | 8 | 4 | 4 | 2 | **1.000** | Additive |
|  | CF00013 | 16 | 8 | 8 | 2 | **0.750** | Additive |
|  | CF00016 | 32 | 16 | 16 | 2 | **0.625** | Additive |
|  | CF00017 | 16 | 4 | 4 | 2 | **0.750** | Additive |
|  | CF00023 | 16 | 4 | 8 | 2 | **0.500** | Synergistic |
|  | CF00038 | 16 | 4 | 4 | 2 | **0.750** | Additive |
|  | CF00040 | 16 | 4 | 4 | 1 | **0.500** | Synergistic |
|  | CF00041 | 16 | 4 | 4 | 2 | **0.750** | Additive |
|  | CF00043 | 8 | 2 | 4 | 1 | **0.500** | Synergistic |
|  | CF00136 | 8 | 2 | 8 | 2 | **0.500** | Synergistic |
|  | CF00258 | 16 | 8 | 4 | 2 | **1.000** | Additive |
|  | CF00855 | 8 | 4 | 8 | 2 | **0.750** | Additive |
|  | CF01975 | 8 | 4 | 4 | 1 | **0.750** | Additive |
|  | CF02033 | 16 | 4 | 8 | 2 | **0.500** | Synergistic |
|  | CF02279 | 4 | 2 | 4 | 0.5 | **0.625** | Additive |
|  | CF02319 | 16 | 8 | 4 | 2 | **1.000** | Additive |
|  | CF02486 | 16 | 4 | 8 | 2 | **0.500** | Synergistic |
| ***M. massiliense*** | CF00008 | 16 | 4 | 8 | 2 | **0.500** | Synergistic |
|  | CF00030 | 16 | 4 | 8 | 2 | **0.500** | Synergistic |
|  | CF00035 | 8 | 4 | 8 | 4 | **1.000** | Additive |
|  | CF00042 | 16 | 4 | 4 | 1 | **0.500** | Synergistic |
|  | CF00046 | 8 | 4 | 8 | 2 | **0.750** | Additive |
|  | CF00047 | 8 | 4 | 4 | 1 | **0.750** | Additive |
|  | CF00883 | 4 | 2 | 4 | 2 | **1.000** | Additive |
| ***M.***  ***bolletii*** | CF00020 | 8 | 4 | 8 | 2 | **0.750** | Additive |
|  | CF00113 | 8 | 4 | 4 | 2 | **1.000** | Additive |
|  | CF00868 | 16 | 8 | 8 | 2 | **0.750** | Additive |
|  | CF02061 | 16 | 4 | 8 | 2 | **0.500** | Synergistic |
| **MIC_50_** | | 16 | 4 | 5.7 | 2 | — | — |
| **MIC_90_** | | 16 | 8 | 8 | 2 | — | — |

**Supplementary Table 7**: Minimum inhibitory concentrations (MICs) of azithromycin and imipenem/relebactam (IMI/REL), both individually and in combination against the *M. abscessus* CF clinical isolates, along with the corresponding fractional inhibitory concentration (FIC) index values and their synergism

| **WGS ID** | **CF Patient ID** | **MIC Value (µg/mL)** | | | | **FIC Index** | **Interaction** |
| --- | --- | --- | --- | --- | --- | --- | --- |
|  |  | **Azithromycin**  **Alone** | **Azithromycin**  **with IMI/REL** | **IMI/REL**  **Alone** | **IMI/REL with Azithromycin** |  |  |
| ***M.***  ***abscessus*** | CF00006 | 4 | 1 | 16 | 2 | **0.375** | Synergistic |
|  | CF00013 | 2 | 1 | 8 | 1 | **0.625** | Additive |
|  | CF00016 | 16 | 8 | 16 | 4 | **0.750** | Additive |
|  | CF00017 | 2 | 1 | 16 | 2 | **0.625** | Additive |
|  | CF00023 | 16 | 4 | 16 | 4 | **0.500** | Synergistic |
|  | CF00038 | 2 | 0.5 | 8 | 1 | **0.375** | Synergistic |
|  | CF00040 | 8 | 2 | 8 | 1 | **0.375** | Synergistic |
|  | CF00041 | 4 | 2 | 4 | 0.5 | **0.625** | Additive |
|  | CF00043 | 8 | 4 | 4 | 1 | **0.750** | Additive |
|  | CF00136 | 4 | 2 | 4 | 1 | **0.750** | Additive |
|  | CF00258 | 16 | 4 | 16 | 4 | **0.500** | Synergistic |
|  | CF00855 | 16 | 4 | 8 | 2 | **0.500** | Synergistic |
|  | CF01975 | 16 | 4 | 16 | 4 | **0.500** | Synergistic |
|  | CF02033 | 16 | 8 | 16 | 4 | **0.750** | Additive |
|  | CF02279 | 32 | 16 | 8 | 2 | **0.750** | Additive |
|  | CF02319 | 8 | 2 | 8 | 4 | **0.750** | Additive |
|  | CF02486 | 4 | 2 | 8 | 2 | **0.750** | Additive |
| ***M. massiliense*** | CF00008 | 4 | 1 | 16 | 0.5 | **0.281** | Synergistic |
|  | CF00030 | 2 | 0.5 | 8 | 1 | **0.375** | Synergistic |
|  | CF00035 | 2 | 0.5 | 8 | 1 | **0.375** | Synergistic |
|  | CF00042 | 4 | 1 | 8 | 2 | **0.500** | Synergistic |
|  | CF00046 | 2 | 1 | 16 | 4 | **0.750** | Additive |
|  | CF00047 | 4 | 2 | 4 | 1 | **0.750** | Additive |
|  | CF00883 | 32 | 16 | 4 | 2 | **1.000** | Additive |
| ***M.***  ***bolletii*** | CF00020 | 32 | 16 | 16 | 8 | **1.000** | Additive |
|  | CF00113 | 32 | 4 | 8 | 4 | **0.625** | Additive |
|  | CF00868 | 16 | 8 | 8 | 4 | **1.000** | Additive |
|  | CF02061 | 8 | 2 | 8 | 2 | **0.500** | Synergistic |
| **MIC_50_** | | 8 | 2 | 8 | 2 | — | — |
| **MIC_90_** | | 32 | 9.8 | 16 | 4 | — | — |

**Supplementary Table 8**: Minimum inhibitory concentrations (MICs) of rifabutin and imipenem/relebactam (IMI/REL), both individually and in combination against the *M. abscessus* CF clinical isolates, along with the corresponding fractional inhibitory concentration (FIC) index values and their synergism

| **WGS ID** | **CF Patient ID** | **MIC Value (µg/mL)** | | | | **FIC Index** | **Interaction** |
| --- | --- | --- | --- | --- | --- | --- | --- |
|  |  | **Rifabutin**  **Alone** | **Rifabutin**  **with IMI/REL** | **IMI/REL**  **Alone** | **IMI/REL with Rifabutin** |  |  |
| ***M.***  ***abscessus*** | CF00006 | 16 | 8 | 16 | 2 | **0.625** | Additive |
|  | CF00013 | 4 | 2 | 4 | 0.5 | **0.625** | Additive |
|  | CF00016 | 16 | 16 | 16 | 0.125 | **1.008** | Indifferent |
|  | CF00017 | 4 | 0.5 | 4 | 2 | **0.625** | Additive |
|  | CF00023 | 16 | 8 | 4 | 1 | **0.750** | Additive |
|  | CF00038 | 16 | 4 | 4 | 1 | **0.500** | Synergistic |
|  | CF00040 | 16 | 2 | 4 | 2 | **0.625** | Additive |
|  | CF00041 | 8 | 1 | 4 | 2 | **0.625** | Additive |
|  | CF00043 | 16 | 4 | 4 | 1 | **0.500** | Synergistic |
|  | CF00136 | 8 | 2 | 16 | 0.125 | **0.258** | Synergistic |
|  | CF00258 | 16 | 4 | 16 | 2 | **0.375** | Synergistic |
|  | CF00855 | 4 | 1 | 4 | 1 | **0.500** | Synergistic |
|  | CF01975 | 2 | 1 | 8 | 0.5 | **0.563** | Additive |
|  | CF02033 | 8 | 4 | 4 | 0.25 | **0.563** | Additive |
|  | CF02279 | 8 | 4 | 2 | 1 | **1.000** | Additive |
|  | CF02319 | 8 | 2 | 4 | 1 | **0.500** | Synergistic |
|  | CF02486 | 16 | 1 | 4 | 2 | **0.563** | Additive |
| ***M. massiliense*** | CF00008 | 16 | 4 | 4 | 2 | **0.750** | Additive |
|  | CF00030 | 16 | 4 | 8 | 2 | **0.500** | Synergistic |
|  | CF00035 | 4 | 1 | 8 | 2 | **0.500** | Synergistic |
|  | CF00042 | 8 | 2 | 4 | 1 | **0.500** | Synergistic |
|  | CF00046 | 16 | 1 | 4 | 2 | **0.563** | Additive |
|  | CF00047 | 8 | 2 | 8 | 2 | **0.500** | Synergistic |
|  | CF00883 | 2 | 0.5 | 4 | 1 | **0.500** | Synergistic |
| ***M.***  ***bolletii*** | CF00020 | 2 | 0.5 | 8 | 1 | **0.375** | Synergistic |
|  | CF00113 | 16 | 4 | 4 | 1 | **0.500** | Synergistic |
|  | CF00868 | 8 | 4 | 4 | 0.5 | **0.625** | Additive |
|  | CF02061 | 8 | 4 | 4 | 0.5 | **0.625** | Additive |
| **MIC_50_** | | 8 | 2 | 4 | 1 | — | — |
| **MIC_90_** | | 16 | 4.9 | 16 | 2 | — | — |

**Supplementary Table 9**: Minimum inhibitory concentrations (MICs) of clofazimine and imipenem/relebactam (IMI/REL), both individually and in combination against the *M. abscessus* CF clinical isolates, along with the corresponding fractional inhibitory concentration (FIC) index values and their synergism

| **WGS ID** | **CF Patient ID** | **MIC Value (µg/mL)** | | | | **FIC Index** | **Interaction** |
| --- | --- | --- | --- | --- | --- | --- | --- |
|  |  | **Clofazimine**  **Alone** | **Clofazimine**  **with IMI/REL** | **IMI/REL**  **Alone** | **IMI/REL with Clofazimine** |  |  |
| ***M.***  ***abscessus*** | CF00006 | 0.5 | 0.5 | 4 | 0.25 | **1.063** | Indifferent |
|  | CF00013 | 0.5 | 0.25 | 2 | 0.25 | **0.625** | Additive |
|  | CF00016 | 0.5 | 0.5 | 8 | 0.25 | **1.031** | Indifferent |
|  | CF00017 | 0.5 | 0.5 | 4 | 0.25 | **1.063** | Indifferent |
|  | CF00023 | 2 | 1 | 8 | 4 | **1.000** | Additive |
|  | CF00038 | 1 | 0.25 | 8 | 4 | **0.750** | Additive |
|  | CF00040 | 2 | 0.5 | 8 | 4 | **0.750** | Additive |
|  | CF00041 | 1 | 0.25 | 8 | 0.25 | **0.281** | Synergistic |
|  | CF00043 | 0.5 | 0.25 | 4 | 2 | **1.000** | Additive |
|  | CF00136 | 2 | 1 | 4 | 2 | **1.000** | Additive |
|  | CF00258 | 2 | 1 | 8 | 1 | **0.625** | Additive |
|  | CF00855 | 1 | 0.25 | 8 | 2 | **0.500** | Synergistic |
|  | CF01975 | 0.5 | 0.25 | 16 | 2 | **0.625** | Additive |
|  | CF02033 | 1 | 0.25 | 4 | 1 | **0.500** | Synergistic |
|  | CF02279 | 2 | 0.5 | 8 | 4 | **0.750** | Additive |
|  | CF02319 | 1 | 0.5 | 16 | 4 | **0.750** | Additive |
|  | CF02486 | 1 | 0.25 | 4 | 1 | **0.500** | Synergistic |
| ***M. massiliense*** | CF00008 | 0.5 | 0.25 | 4 | 2 | **1.000** | Additive |
|  | CF00030 | 2 | 1 | 8 | 1 | **0.625** | Additive |
|  | CF00035 | 1 | 0.5 | 16 | 4 | **0.750** | Additive |
|  | CF00042 | 2 | 1 | 8 | 8 | **1.500** | Indifferent |
|  | CF00046 | 2 | 1 | 16 | 8 | **1.000** | Additive |
|  | CF00047 | 2 | 1 | 8 | 1 | **0.625** | Additive |
|  | CF00883 | 1 | 0.5 | 4 | 1 | **0.750** | Additive |
| ***M.***  ***bolletii*** | CF00020 | 1 | 0.5 | 16 | 4 | **0.750** | Additive |
|  | CF00113 | 1 | 0.5 | 8 | 4 | **1.000** | Additive |
|  | CF00868 | 1 | 0.5 | 16 | 4 | **0.750** | Additive |
|  | CF02061 | 1 | 0.5 | 8 | 2 | **0.750** | Additive |
| **MIC_50_** | | 1 | 0.5 | 8 | 2 | — | — |
| **MIC_90_** | | 2 | 1 | 16 | 4 | — | — |

**Supplementary Table 10**: Minimum inhibitory concentrations (MICs) of minocycline and imipenem/relebactam (IMI/REL), both individually and in combination against the *M. abscessus* CF clinical isolates, along with the corresponding fractional inhibitory concentration (FIC) index values and their synergism

| **WGS ID** | **CF Patient ID** | **MIC Value (µg/mL)** | | | | **FIC Index** | **Interaction** |
| --- | --- | --- | --- | --- | --- | --- | --- |
|  |  | **Minocycline**  **Alone** | **Minocycline**  **with IMI/REL** | **IMI/REL**  **Alone** | **IMI/REL with Minocycline** |  |  |
| ***M.***  ***abscessus*** | CF00006 | 256 | 128 | 4 | 2 | **1.000** | Additive |
|  | CF00013 | 256 | 128 | 4 | 1 | **0.750** | Additive |
|  | CF00016 | 512 | 256 | 32 | 8 | **0.750** | Additive |
|  | CF00017 | 256 | 128 | 2 | 2 | **1.500** | Indifferent |
|  | CF00023 | 512 | 64 | 4 | 2 | **0.625** | Additive |
|  | CF00038 | 256 | 64 | 4 | 1 | **0.500** | Synergistic |
|  | CF00040 | 256 | 128 | 8 | 4 | **1.000** | Additive |
|  | CF00041 | 512 | 128 | 2 | 2 | **1.250** | Indifferent |
|  | CF00043 | 256 | 128 | 2 | 2 | **1.500** | Indifferent |
|  | CF00136 | 256 | 64 | 2 | 1 | **0.750** | Additive |
|  | CF00258 | 256 | 128 | 8 | 2 | **0.750** | Additive |
|  | CF00855 | 256 | 128 | 8 | 2 | **0.750** | Additive |
|  | CF01975 | 256 | 64 | 8 | 2 | **0.500** | Synergistic |
|  | CF02033 | 256 | 128 | 4 | 2 | **1.000** | Additive |
|  | CF02279 | 256 | 128 | 2 | 1 | **1.000** | Additive |
|  | CF02319 | 256 | 64 | 2 | 1 | **0.750** | Additive |
|  | CF02486 | 1024 | 128 | 8 | 4 | **0.625** | Additive |
| ***M. massiliense*** | CF00008 | 256 | 64 | 8 | 2 | **0.500** | Synergistic |
|  | CF00030 | 256 | 16 | 4 | 2 | **0.563** | Additive |
|  | CF00035 | 256 | 32 | 4 | 0.5 | **0.250** | Synergistic |
|  | CF00042 | 16 | 16 | 4 | 0.25 | **1.063** | Indifferent |
|  | CF00046 | 512 | 128 | 4 | 1 | **0.500** | Synergistic |
|  | CF00047 | 256 | 32 | 2 | 2 | **1.125** | Indifferent |
|  | CF00883 | 8 | 8 | 2 | 0.125 | **1.063** | Indifferent |
| ***M.***  ***bolletii*** | CF00020 | 256 | 256 | 4 | 2 | **1.500** | Indifferent |
|  | CF00113 | 256 | 64 | 4 | 1 | **0.500** | Synergistic |
|  | CF00868 | 8 | 8 | 4 | 0.125 | **1.031** | Indifferent |
|  | CF02061 | 256 | 64 | 4 | 2 | **0.750** | Additive |
| **MIC_50_** | | 256 | 90.5 | 4 | 2 | — | — |
| **MIC_90_** | | 512 | 128 | 8 | 2.5 | — | — |

**Supplementary Figure 1**: Bacterial load (log_10_ CFU/ml) of *M. abscessu*s ATCC 19977 over 72 hours with single-agent therapies and three-drug combination therapies with imipenem/relebactam (IMI/REL) at 16× MIC, 4× MIC, 1× MIC, and 1/4× MIC

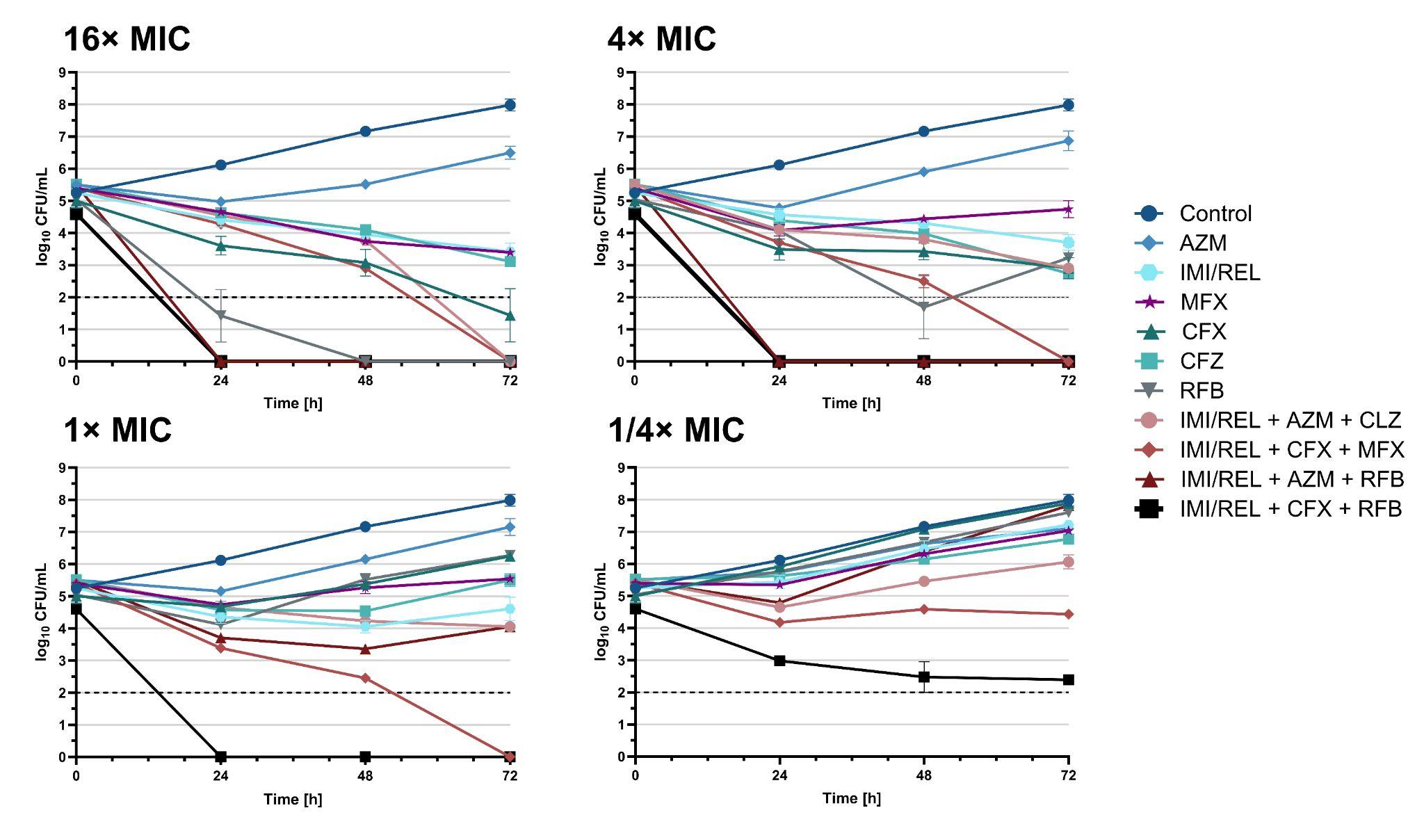

Data are presented as mean with standard errors of the mean. The horizontal dashed line marks the lower limit of detection.

MIC = minimum inhibitory concentration, AZM = azithromycin, MFX = moxifloxacin, CFX = cefoxitin, CFZ = clofazimine, RFB = rifabutin

**Supplementary Figure 2**: Bacterial load (log_10_ CFU/ml) of *M. abscessu*s CF clinical isolate 13 over 72 hours with imipenem/relebactam (IMI/REL), cefoxitin, rifabutin, and their combination at 16× MIC, 4× MIC, 1× MIC, and 1/4× MIC

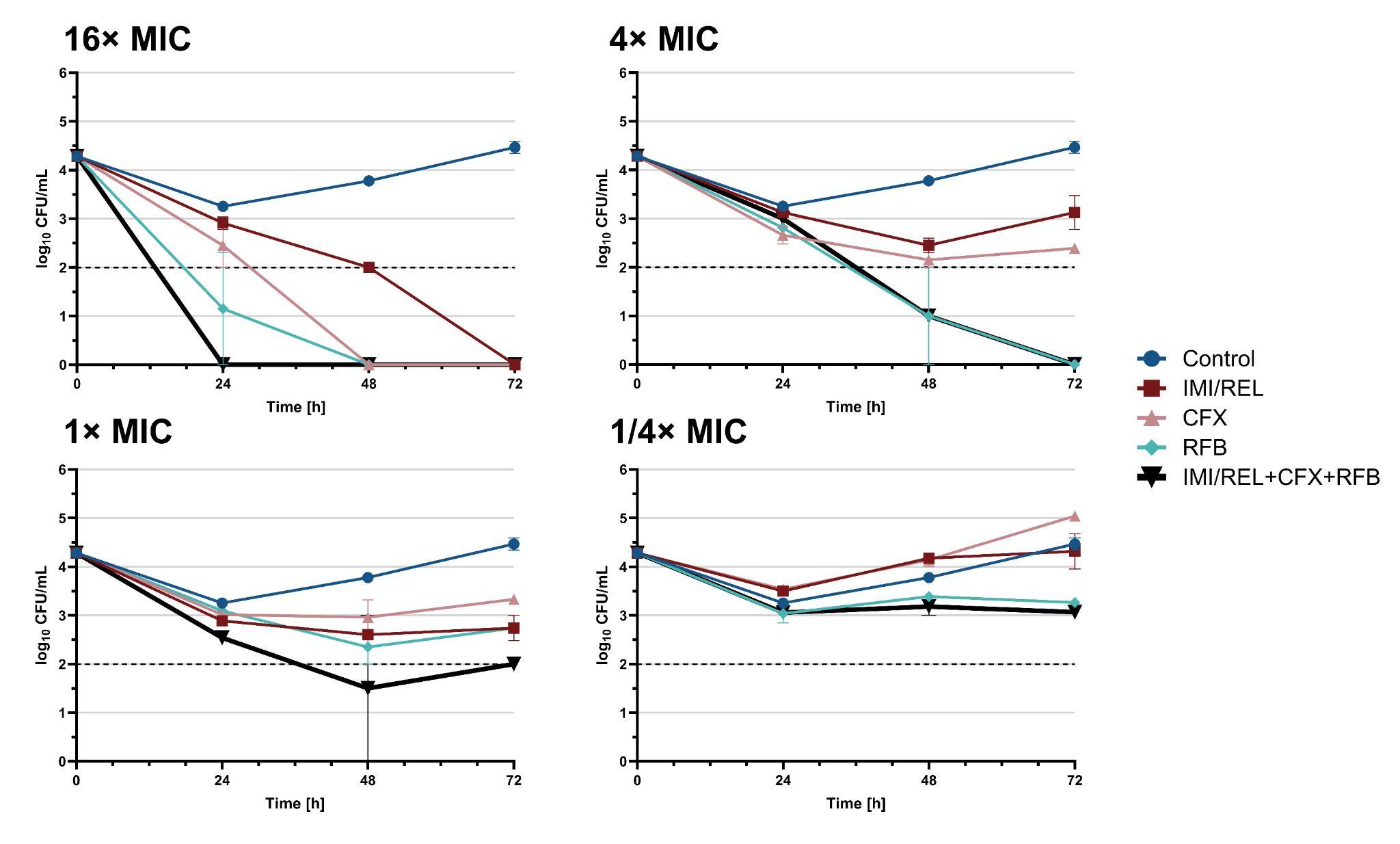

Data are presented as mean with standard errors of the mean. The horizontal dashed line marks the lower limit of detection.

MIC = minimum inhibitory concentration, CFX = cefoxitin, RFB = rifabutin

**Supplementary Figure 3**: Bacterial load (log_10_ CFU/ml) of *M. abscessu*s CF clinical isolate 258 over 72 hours with imipenem/relebactam (IMI/REL), cefoxitin, rifabutin, and their combination at 16× MIC, 4× MIC, 1× MIC, and 1/4× MIC.

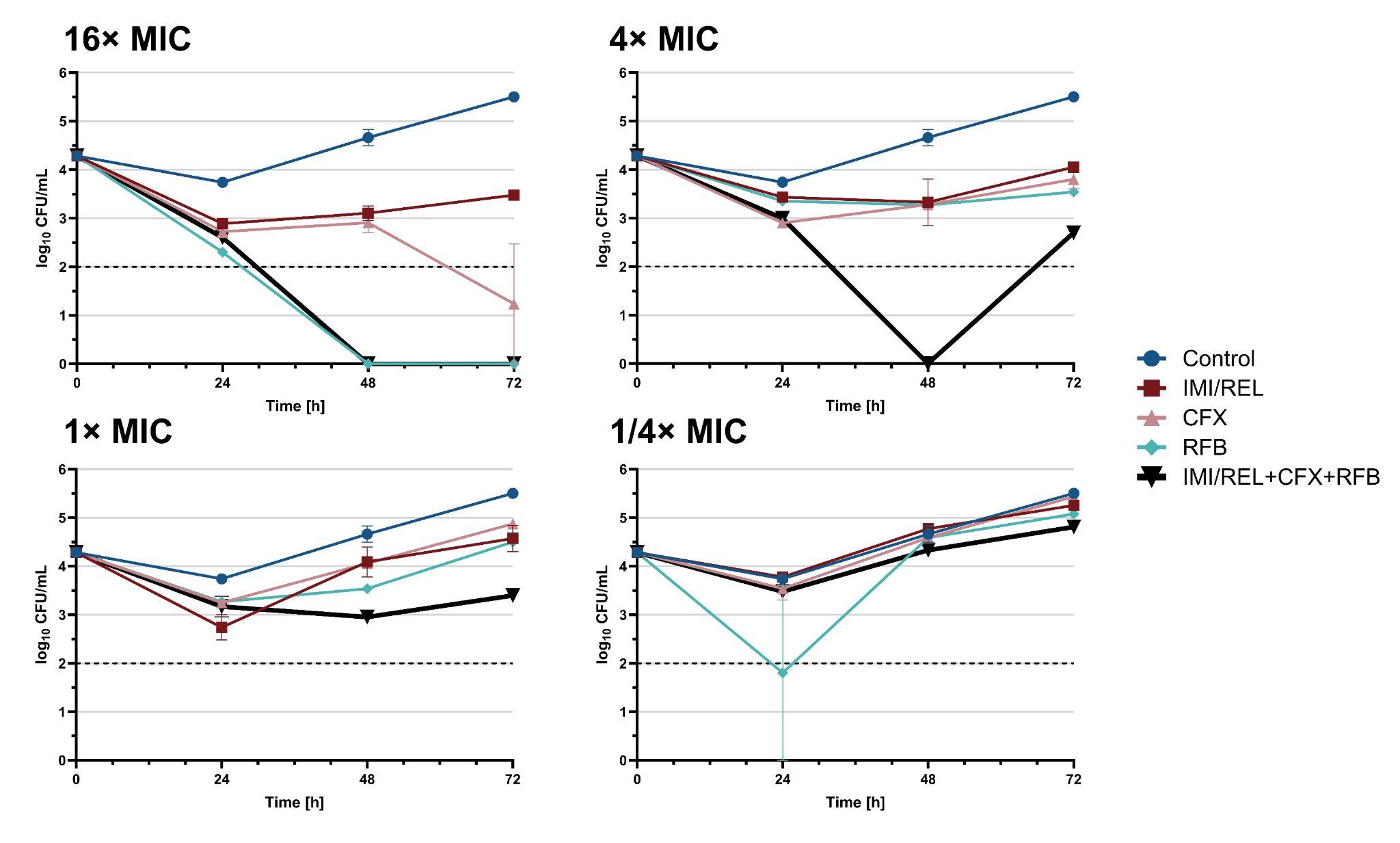

Data are presented as mean with standard errors of the mean. The horizontal dashed line marks the lower limit of detection.

MIC = minimum inhibitory concentration, CFX = cefoxitin, RFB = rifabutin

**Supplementary Table 11**: Minimum inhibitory concentrations (MICs) of imipenem and imipenem/relebactam (IMI/REL) against the *M. abscessus* CF clinical isolates.

| **WGS ID** | **CF Patient ID** | **MIC Value (µg/mL)** | |
| --- | --- | --- | --- |
|  |  | **Imipenem/Relebactam^a^** | **Imipenem** |
| ***M. abscessus*** | ATCC 19977 | 4 | 8 |
|  | CF00006 | 16 | 8 |
|  | CF00013 | 4 | 8 |
|  | CF00016 | 16 | 8 |
|  | CF00017 | 4 | 8 |
|  | CF00023 | 4 | 8 |
|  | CF00038 | 4 | 8 |
|  | CF00040 | 8 | 8 |
|  | CF00041 | 4 | 8 |
|  | CF00043 | 4 | 8 |
|  | CF00136 | 8 | 16 |
|  | CF00258 | 8 | 8 |
|  | CF00855 | 8 | 4 |
|  | CF01975 | 16 | 16 |
|  | CF02033 | 4 | 8 |
|  | CF02279 | 4 | 16 |
|  | CF02319 | 4 | 8 |
|  | CF02486 | 8 | 8 |
| ***M. massiliense*** | CF00008 | 8 | 16 |
|  | CF00030 | 8 | 8 |
|  | CF00035 | 8 | 8 |
|  | CF00042 | 8 | 8 |
|  | CF00046 | 8 | 8 |
|  | CF00047 | 4 | 8 |
|  | CF00883 | 4 | 4 |
| ***M. bolletii*** | CF00020 | 8 | 16 |
|  | CF00113 | 4 | 4 |
|  | CF00868 | 8 | 8 |
|  | CF02061 | 4 | 4 |
| **MIC_50_** | | 8 | 8 |
| **MIC_90_** | | 9.2 | 16 |

^a^ Represents the median MIC value for each strain across all susceptibility assays.
